# A Portable Impedance Microflow Cytometer for Measuring Cellular Response to Hypoxia

**DOI:** 10.1101/2020.07.28.224006

**Authors:** Darryl Dieujuste, Yuhao Qiang, E Du

## Abstract

This paper presents the development and testing of a low-cost, portable microflow cytometer based on electrical impedance sensing, for single cell analysis under controlled oxygen microenvironment. The cytometer system is based on an AD5933 impedance analyzer chip, a microfluidic chip, and an Arduino microcontroller operated by a custom Android application. A representative case study on human red blood cells (RBCs) affected by sickle cell disease is conducted to demonstrate the capability of the cytometry system. Equivalent circuit model of a suspending biological cell is used to interpret the electrical impedance of single flowing RBCs. In normal blood, cytoplasmic resistance and membrane capacitance do not change significantly with the change in oxygen tension. In contrast, RBCs affected by sickle cell disease show that upon hypoxia treatment, the cytoplasmic resistance decrease from 11.6 MΩ to 23.4 MΩ, and membrane capacitance decrease from 1.1 pF to 0.8 pF. Strong correlations are identified between the changes in these subcellular electrical components of single cells and the cell sickling process induced by hypoxia treatment. The representative results reported in this paper suggest that single cell electrical impedance can be used as a sensitive biophysical marker for quantifying cellular response to change in oxygen concentration. The developed flow cytometry system and the methodology can also be extended to analysis of cellular response to hypoxia in other cell types.

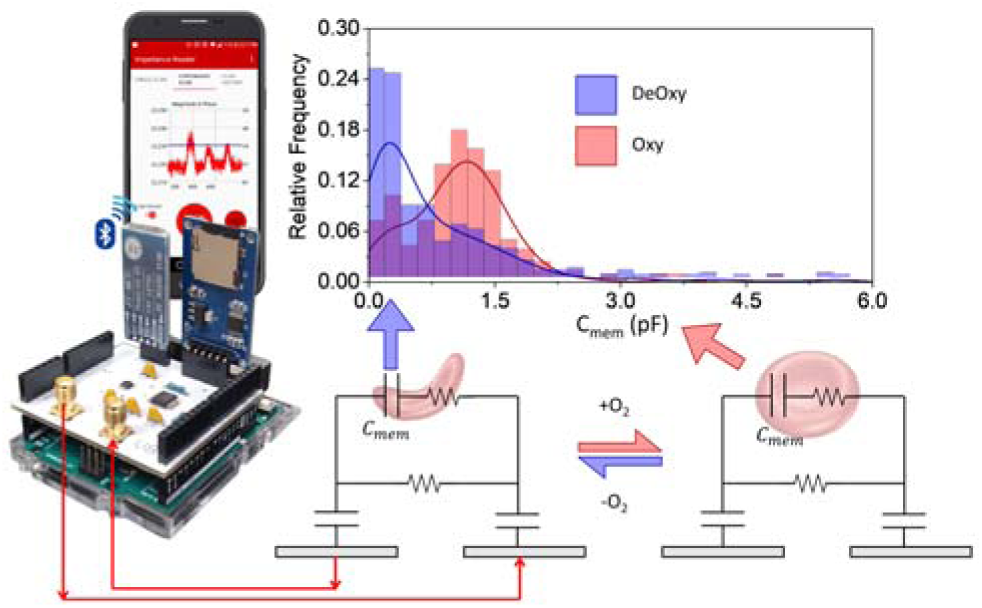

## I. Introduction

Hypoxia (deprivation of oxygen in the body) causes a variety of physiological changes in cells. Physiological responses to hypoxia due to high-altitude or deep-sea diving, or pathological responses have been extensively investigated at the whole-body level and at the single cell levels [1, 2]. Analysis of single-cell suspensions has become of important medical interest. Study of cellular responses to hypoxia has provided insights in tumor pathology [3], cancer treatment [4], cardiovascular pathophysiology [5], metabolism [6, 7], and homeostatic mechanism in mammalian cells [8]. A golden standard to measure cellular hypoxia and response to hypoxia environment is via flow cytometry analysis of single cells by measuring the protein levels, such as hypoxia-inducible factor 1-alpha (HIF1α) and BCL2/adenovirus E1B 19 kDa protein interacting protein 3 (BNIP3) [9, 10]. This method provides high specificity via antibody-based immunostaining to target the proteins of interest, but also requires the analyzed cells being fixed and permeabilized.

Recently, electrical impedance-based flow cytometry has been demonstrated as an alternative method to the conventional optical approach in the analysis of single cells. It is inherently quantitative, non-invasive and label-free, eliminating the needs for fluorescence or biochemical labeling [11]. Electrical impedance-based flow cytometry of single cells has been demonstrated to be useful to discriminate red blood cell (RBC) that are affected by malaria [12, 13] and sickle cell disease (SCD) [14]. In the study of cellular response to hypoxia, microfluidics technology can be used to precisely control the gaseous microenvironment and allow simultaneous microscopy and impedance sensing of cells in flow and in stationary conditions [15]. However, these electrical impedance analyses were performed using large benchtop equipment. Miniaturization of electrical-impedance based flow cytometry systems can provide field-testing and point-of-care assay of single cells in response to hypoxia. Portable electrical impedance systems have been created for general bio-impedance purposes [16]. Microcontrollers with peripheral devices for user control and observation have been used with the popular AD5933 integrated circuit [17] to create portable impedance sensors [18, 19]. Further portability is achieved when implementing wireless communication such as Bluetooth to separate data acquisition on the sensor from data processing on the personal computer (PC) [20]. Control of the sensor and visualization of the data can be managed using smart phone applications via wireless communication [21].

In this work, we present the development and testing of a portable, mobile app controlled, impedance-based flow cytometer for measuring cellular response to hypoxia. We demonstrate the capability of electrical impedance-based flow cytometry of single cells using red blood cells (RBCs), as they are the most abundant type of cells and contain oxygen carrier protein, hemoglobin (Hb), which is sensitive to the environmental oxygen tension. We show discrimination between normoxic and hypoxic states for the same population of RBCs. We further demonstrate that this portable device is sensitive to discriminate RBCs from healthy donor and SCD patients. In SCD, phenotypic response of the mutation of normal HbA into HbS is the formations of rigid HbS fibers and sickled shape as the HbS polymerize in response to low oxygen tension (hypoxia) as opposed to oxygenated condition (normoxia) [22, 23]. The portable system can be readily used as a point-of-care diagnosis and monitoring tool of SCD, which is conventionally diagnosed in labs via techniques for detection of HbS, such as hemoglobin electrophoresis [24], isoelectric focusing [25] or high-performance liquid chromatography [26], chromatographic immunoassay [27], and density-based erythrocyte measurement [28].

## II. Materials and methods

### A. Microfluidic Chip Design

The electrical impedance measurement of RBCs was carried out on a microfluidic chip (Figure 1A). The microfluidic chip is a set of microchannel and two parallel Ti/Au electrodes (with 20 μm gap and 20 μm band width) coated on a 0.7 mm thick glass. The microchannel consists two layers of polydimethylsiloxane (PDMS) channel: the top layer is a serpentine shape gas channel and the bottom one is a thin gas permeable membrane (150 μm thick) with a straight channel for cell suspension. Both PDMS channels were fabricated following a standard soft lithography method. The narrowest portion of the microchannel measures 20 μm wide and 5 μm deep. Electrodes were deposited and patterned on a glass substrate using E-beam evaporation and standard photolithography techniques [29]. The double-layer microchannel and electrodes-glass substrate are bonded together using air plasma. Conductive wires were soldered to the electrodes to allow the cells to be measured by the portable impedance sensing system.

**Figure 1.**
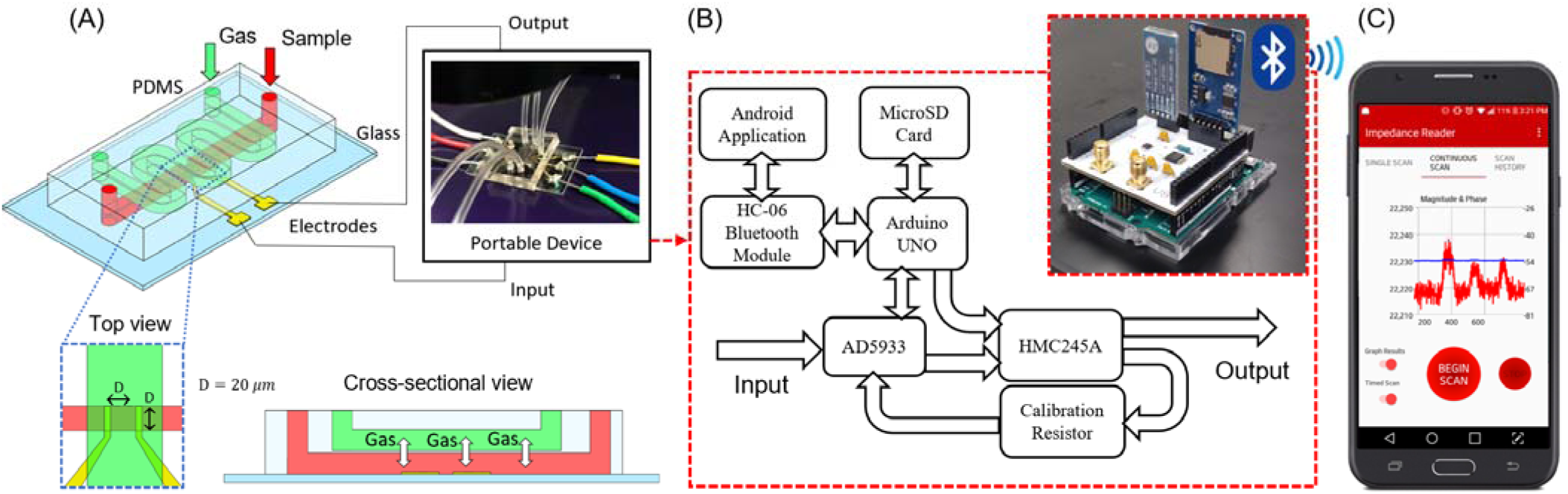
Device overview of the portable impedance-based flow cytometer prototype. (A) A double layer PDMS device bonded to a glass substrate patterned with Ti/Au electrodes. The electrodes provide connectivity to the portable device. The top view shows the intersection of the gas channel over the point of measurement between the electrodes. The cross-sectional view depicts the gas exchange between two channels to induce cell sickling. (B) The flow chart of the portable device consisting of all major components used and how information moves between the components. (C) The Android application used to control the portable device operated to continuously scan for a designated length of time and produce a graph of the results.

### B. Portable impedance-based flow cytometer prototype design

Figure 1B illustrates the impedance sensing system as used during experimentation for validation. The prototype is centered around the AD5933 impedance converter by Analog Devices, for its ability to output a sinusoidal wave with controlled frequency and voltage as well as calculating and providing the admittance of an unknown sample. A printed circuit board (PCB) was designed to connect directly to a commercial Arduino UNO microcontroller. On the PCB, the AD5933 is connected to the Arduino UNO using the two-wire I2C communication protocol. The Arduino UNO is operated via an Android application through Bluetooth communication using an HC-06 Bluetooth Module. The Bluetooth module uses a two-wire serial connection to relay data to and from the Arduino UNO. The Android application allows the user to begin impedimetric scans over a set duration or until manually stopped. Scan results can be optionally plotted on phone and a text file containing the numerical data can be distributed via email. The Arduino UNO and the AD5933 are connected to an HMC245A switch. The switch provides calibration functionality for the AD5933 by establishing a multi output, single input signal path. An SD card reader is connected to the Arduino UNO using a 3-wire SPI connection. The SD card provides large storage capabilities for fast data throughput needed for flow cytometry in addition to providing an easy means of data transfer to PCs for further analysis. Two female SMA connectors on the PCB provide connectivity to the soldered wires on the microfluidic chip.

### C. Experimental Protocol

The inlet to the gas channel is inserted with a standard 0.02 in. ID/0.06 in. OD microbore tubing and connected to a PC controlled switching valve via to alternate between a high purity N_2_ gas and a gas mixture of 5% CO_2_, 17.5% O_2_ and 77.5% N_2_. For each patient sample, 5 µL of cell suspension was diluted into 1 mL of phosphate buffered saline (PBS). The cell channel was primed with PBS. Microliter samples of RBC suspension are injected into the suspension channel via the microbore tubing through 1.03 mm, 50 µL, Hamilton glass syringe. The flow rate of cell suspension is fixed at 100 pL/min using a Harvard Apparatus Pump 11 Pico Plus Elite syringe pump. The portable cytometer is operated to measure data continuously using the custom Android application. An example of the application interface displaying results is shown in Figure 1C. Simultaneous microscope video is recorded in order to validate impedimetric readings. Data is saved onto the SD card during the experiment and analyzed by a custom MATLAB script.

### D. Statistical Analysis

All data are expressed as mean ± SD. Statistical analyses of RBCs were performed with OriginPro 2020 (OriginLab). Mann Whitney test between measurements of samples under Oxy and DeOxy was used to generate the *p* values. A p-value less than 0.05 is statistically significant. Sample distribution was fitted based on the Kernel density estimation.

## III. Results and discussion

### A. Portable Device Data Analysis

The impedance response of individual RBCs on the portable device was validated by comparing it with the response of cells measured using the HF2IS Impedance Spectroscope by Zurich Instruments. Results obtained using the portable device, after being filtered in MATLAB, return impedance values of comparable magnitude and resolution to the benchtop equipment. Figure 2 displays impedance of an RBC, in the form of bell-shaped curves, measured by a bench top HF2IS instrument and by our portable device. The curve represents a cell passing through the pair of electrodes compared to the impedance of PBS medium only. A level 3 wavelet filter with BlockJS denoising in MATLAB was used to denoise the raw signal (gray color, Figure 2C) obtained from the portable device, which produces impedance measurement comparable to the commercial instrument. MATLAB algorithm was used to analyze the magnitude of the impedance which allowed the detection of peaks as shown in Figure 3. The peak location is then stored and used to collect the other part of impedance information (phase) from the original data.

**Figure 2.**
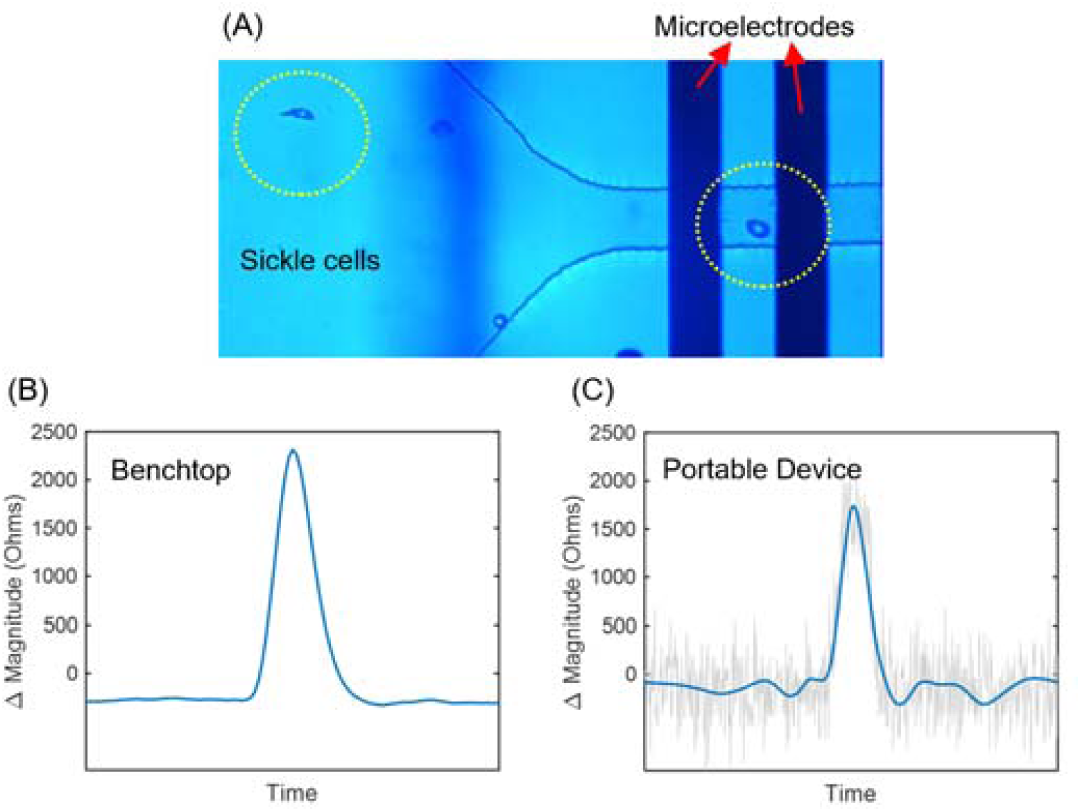
(A) Microscope image of RBCs under hypoxia travelling through the microfluidic channel to be measured between two microelectrodes. (B) The impedance magnitude response of a single cell passing through the microelectrode pair using the HF2IS. (C) The impedance magnitude response of a single cell passing through the microelectrode pair using the portable device. The gray signal is the raw response recorded. The blue signal is result of passing the raw response through a level 3 wavelet filter with BlockJS denoising in MATLAB.

**Figure 3.**
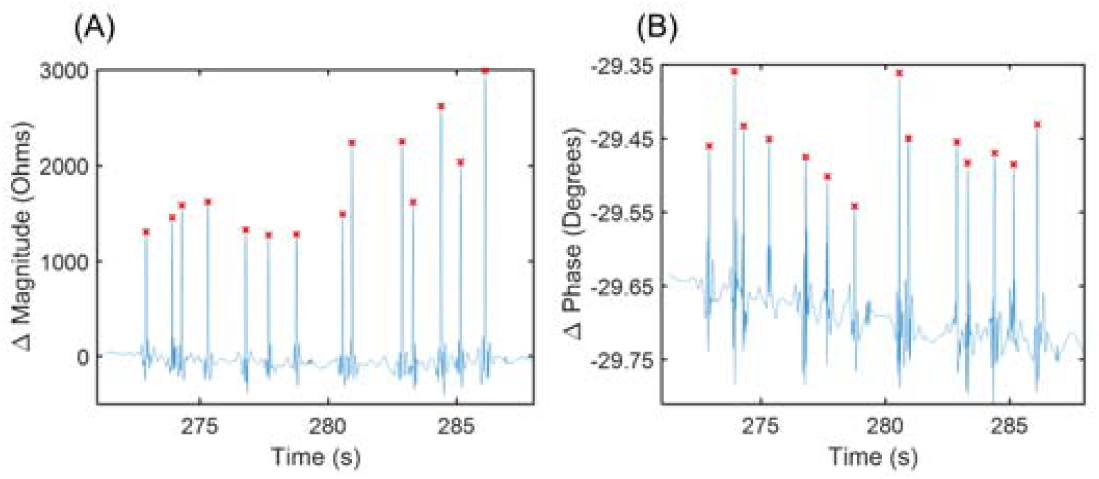
(A) A MATLAB algorithm is used to identify fourteen peaks from a detrended impedance measurement (magnitude results) obtained from the portable device using a minimum threshold. (B) Corresponding phase values were obtained by matching the time coordinate from the magnitude plot.

Impedance of a single cell is calculated utilizing the equivalent circuit models as shown in Figure 4. In the case of PBS medium only, the equivalent circuit model consists of medium impedance, *z*_*PBS*_ in series of an electric double layer impedance, simplified to a single variable, *z*_*dlt*_, the total impedance is,

**Figure 4.**
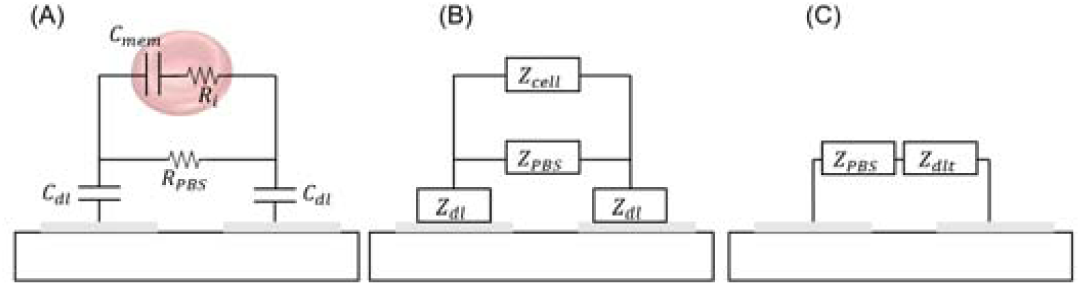
(A) An electrical circuit model of an RBC in a PBS medium. R_l_ and C_mem_ represent the internal resistance of the cell and the capacitance of the cell membrane respectively. C_dl_ is the double layer capacitance created between the electrodes and PBS medium with a resistance, R_PBS_. (B) The components in part A are simplified to represent impedances. (C) The circuit shown represents the measured impedance when a cell is not present. Z_dlt_: total double layer impedance from both electrodes.

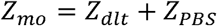

where *z*_*dlt*_ is purely capacitive and *z*_*PBS*_ purely resistive. Under these assumptions, the real part of the cartesian representation of *z*_*mo*_ will be equal to *z*_*PBS*_ and the imaginary part is equal to *z*_*dlt*_.

When a cell is present, the total impedance measured becomes,

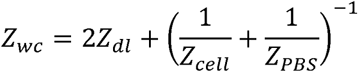

The individual cell impedance can then be calculated by,

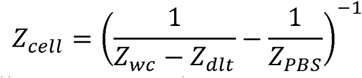

The impedance for each measured cell was converted to a cartesian equivalent. The real part of the cartesian impedance is assumed to reflect the internal resistance, *R*_*i*_, of the cell and the imaginary part is assumed to reflect the membrane capacitance. Value of membrane capacitance is then calculated by

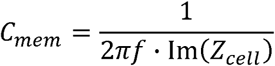

where *f* is the frequency used for the impedimetric measurement. The calculated membrane capacitance is plotted against its respective internal resistance for each cell.

### B. Normal RBC and SS RBC Results

Figure 5 groups the measured impedance values RBCs from two normal blood samples (AA1 and AA2) and sickle cell samples from three SCD patients (SS1-SS3). For normal RBCs, the distributions in both the relative magnitude and relative phase show remarkable overlap between Oxy and DeOxy conditions (Figures 5 AB). This suggests that hypoxia treatment does not affect electrical impedance of single normal RBCs significantly. For SS RBCs, the relative magnitude and relative phase were more spread than normal cells, disregarding the level of oxygen tension. This was not surprising as SS RBCs are more heterogeneous in terms of cell volume and other intrinsic properties, such as membrane damage and intracellular Hb variants. In addition, SS samples show significant change in their distribution upon hypoxia treatment in terms of both magnitude and phase, from a weak bimodality in Oxy condition to a single modality of smaller mean value in DeOxy condition (Figures 5 CD). This could be explained by the cell sickling process along with intracellular HbS polymerization, resulting in reduced cell volume and a denser distribution of intracellular variations. Table 1 provides the mean and standard deviation values for each cell condition.

**Table 1.**
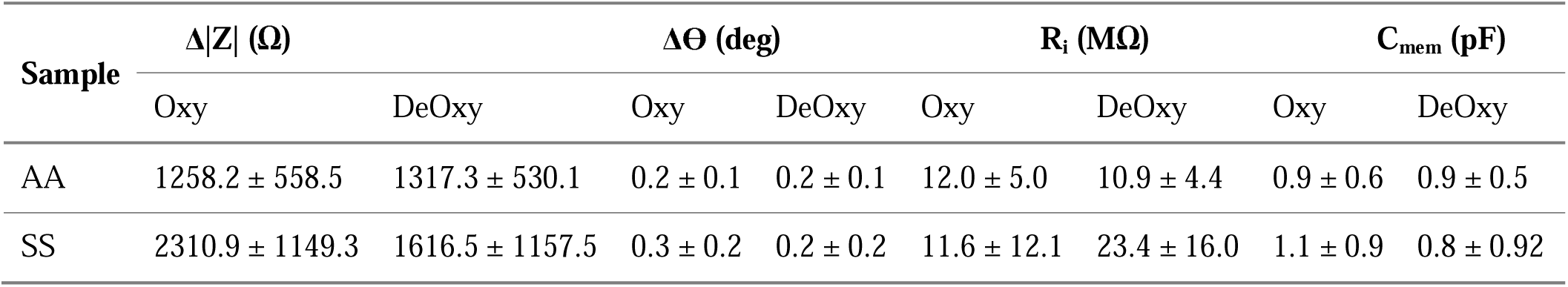
Electrical impedance and single cell electrical properties of normal and SS RBCs. AA-Oxy (n = 162), AA-DeOxy (n = 267), SS-Oxy (n = 274), SS-DeOxy (n = 343).

| Sample | $\Delta\vert Z\vert $ ($\Omega$) | | $\Delta\Theta$ (deg) | | $R_i$ ($M\Omega$) | | $C_{mem}$ (pF) | |
| --- | --- | --- | --- | --- | --- | --- | --- | --- |
|  | Oxy | DeOxy | Oxy | DeOxy | Oxy | DeOxy | Oxy | DeOxy |
| AA | $1258.2 \pm 558.5$ | $1317.3 \pm 530.1$ | $0.2 \pm 0.1$ | $0.2 \pm 0.1$ | $12.0 \pm 5.0$ | $10.9 \pm 4.4$ | $0.9 \pm 0.6$ | $0.9 \pm 0.5$ |
| SS | $2310.9 \pm 1149.3$ | $1616.5 \pm 1157.5$ | $0.3 \pm 0.2$ | $0.2 \pm 0.2$ | $11.6 \pm 12.1$ | $23.4 \pm 16.0$ | $1.1 \pm 0.9$ | $0.8 \pm 0.92$ |

**Figure 5.**
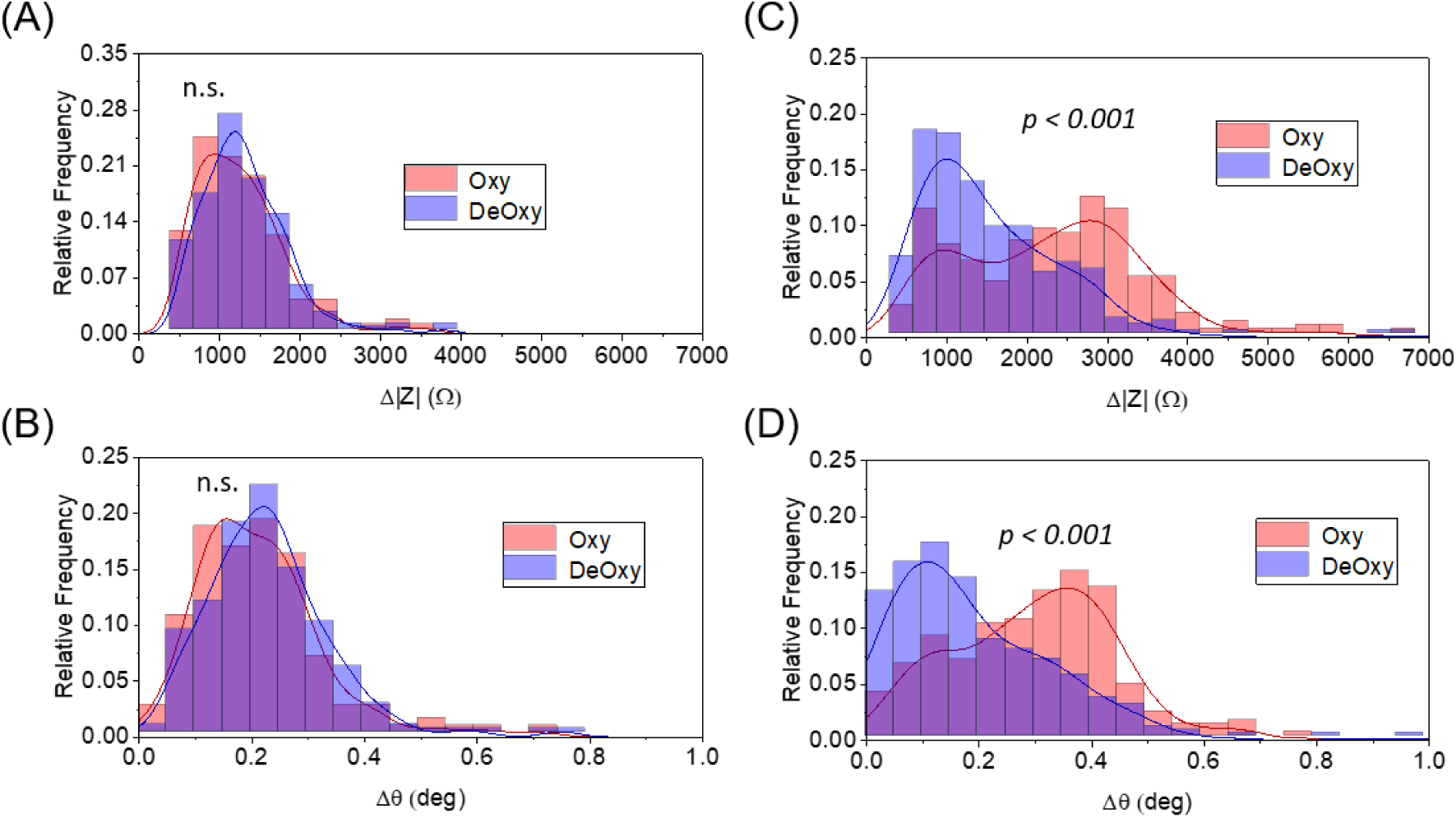
(A) Histogram depicting the relative magnitude of the impedance for detected healthy cells in oxygenated (Oxy) and deoxygenated (DeOxy) conditions. The data includes samples AA1 and AA2. (B) The corresponding relative phase of the impedance for detected healthy cells from A. (C) The relative magnitude of the impedance for detected sickle cells in Oxy and DeOxy conditions. The data includes samples SS1, SS2, and SS3. (D) The corresponding relative phase of the impedance for detected sickle cells in C. n.s. stands for not significant.

To verify the contributions from the subcellular components, i.e. membrane and cytoplasm upon hypoxia to the single cell electrical impedance, we calculated the membrane capacitance and internal resistance for each cell group, using equivalent circuit models described in Figure 4A. Table 1 provides the mean and standard deviation values of single cell analysis results for both normal and SS RBCs under Oxy and DeOxy conditions. Figure 6 shows the distribution of single cell analysis. Similar to the impedance results, there is not a significant change in either parameter for normal cells transitioning from an Oxy to DeOxy state. However, we observed significant changes in both values for SCD samples.

**Figure 6.**
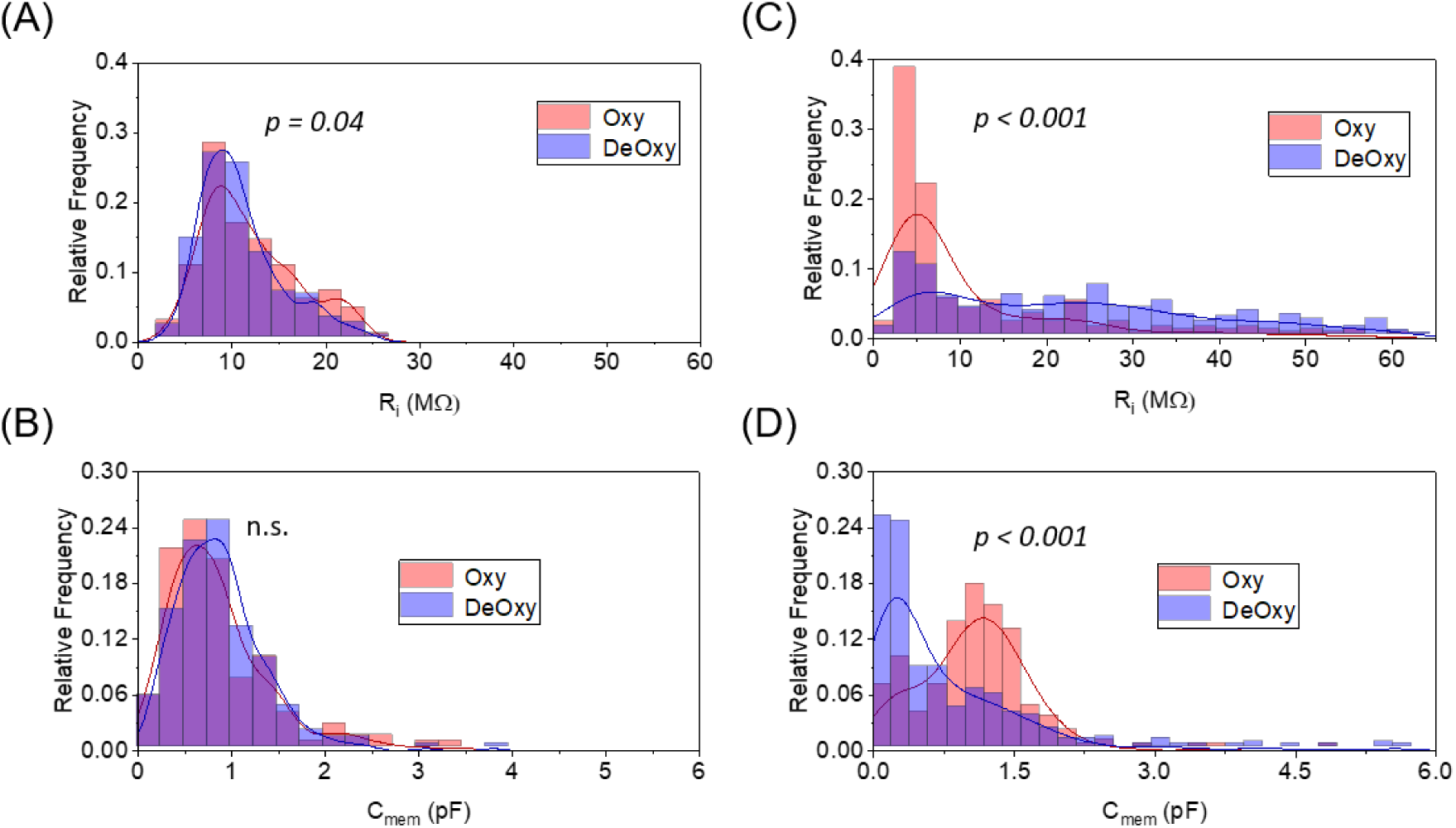
(A) This graph depicts the calculated internal resistance for detected healthy cells in oxygenated (Oxy) and deoxygenated (DeOxy) conditions. The data includes samples AA1 and AA2. (B) The calculated membrane capacitance for detected healthy cells from A. (C) The calculated internal resistance for detected sickle cells in Oxy and DeOxy conditions. The data includes samples SS1, SS2, and SS3. (D) The calculated membrane capacitance for detected sickle cells from C. n.s. stands for not significant.

### C. Discussion

The changes recorded from normoxia to hypoxia vary from sample to sample, but the collective trends observed when grouping samples provide a means of understanding the measured data. We attempt to understand the changes in cellular internal resistance and membrane capacitance by correlating the values to the physical changes occurring during cell sickling. Generally, variations in subcellular electric components may be interpreted using a simple relationship between resistance, electrical resistivity (ρ), length (L) and cross-sectional area (A) of a specimen using the following equation

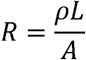

The lipid bilayer of a cell membrane is typically viewed as a parallel-plate capacitor. Capacitance of a standard parallel-plate capacitor can be defined as a function of a specimen’s dielectric permittivity (□), cross-sectional area (A), and distance between the plates (d),

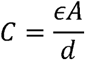

These relations can be applied to a biological cell and we can say the cell’s cross-sectional area is directly proportional to the membrane capacitance and inversely proportional to the internal resistance. During cell sickling, the cross-sectional area of a sickled cell can be reduced significantly if the cell takes on its classical sickle shape. Normal RBCs are not expected to exhibit significant changes to their morphology and therefore the cross-sectional area of these cells should remain fairly consistent. This is supported by the trend of results in Figure 6.

It should also be noted that single cell analysis of normal RBCs showed a significant negative shift in internal resistance (*p = 0*.*04*, Figure 6A). This may be associated with a change in the water-protein structure in aqueous solution[30], which makes Oxy Hb slightly more resistive to electric currents than DeOxy Hb. In contrast, for SS RBCs, single cell analysis showed the significant positive shift in the mean internal resistance with the value almost doubled upon hypoxia treatment. Interestingly, the internal resistance showed a remarkably wider distribution upon hypoxia, indicating the marked heterogeneity in HbS polymerization and the consequent effective subcellular resistance to current. The significant changes in both subcellular electric components for SS RBCs are largely attributed to the polymerization process of intracellular HbS, different from the mechanism for normal RBCs.

## IV. Conclusion

Impedance-based flow cytometry is a label-free, non-invasive method for cell analysis. We have utilized this technique in our portable, mobile app-controlled device in tandem with our PDMS based microfluidic chip to perform an impedimetric analysis of cell samples under normoxia and hypoxia conditions. The experimental results are positively correlated with mathematical models to provide probable explanations to observed trends. There is potential for utilizing this work for as a foundation for a diagnosis method for SCD. The changes in the single cell electrical impedance can serve as a potential biophysical marker for SCD. The developed single cell analysis system can be extended to study cellular response to hypoxia for other cell types.

## V. Acknowledgement

This work was supported by the NSF Grant No. 1635312 and NIH Grant 1OT2HL152638. D.D. acknowledges the support from NIH R01EB025819-03S1. E.D. and Y.Q. acknowledge support from NIH Grant 5R01EB025819. The authors thank Dr. Ofelia Alvarez at the Division of Pediatric Hematology and Oncology, University of Miami for providing sickle cell blood samples. D.D. would like to thank Dr. Jia Liu for discussion on the microfluidic setup.

